# A Multi-Modal Genomic Knowledge Distillation Framework for Drug Response Prediction

**DOI:** 10.1101/2024.10.17.618828

**Authors:** Shuang Ge, Shuqing Sun, Huan Xu, Qiang Cheng, Zhixiang Ren

## Abstract

Precision oncology utilizes genomic data to tailor treatment to individuals. Cancer drug sensitivity studies can predict the response levels of different drugs for the same cultured cancer cell line, which is beneficial for personalized medicine. Recent studies have demonstrated that integrating multi-modal genomic data, e.g., gene expression, mutation, copy number alteration, methylation, can provide comprehensive knowledge and improve drug response prediction. Although multimodal genomic profiles are generally available from public datasets, only gene expression data is commonly used in clinical settings. In this study, we propose a framework for privileged information knowledge distillation to transfer knowledge from a multi-modal genomic teacher network, using only gene expression for inference. Specifically, we train a teacher network by feature re-weighting based on inter-modality dependencies and align the inter-sample correlations through our proposed relation-aware differentiation distillation. Experiments on the Genomics of Drug Sensitivity in Cancer (GDSC) dataset demonstrate that our framework improves drug response prediction by about 6% compared to the baseline and outperforms state-of-the-art methods. Transferable studies performed on missing GDSC data and clinical datasets further confirm the feasibility of our model for predicting drug responses using only gene expression data.

## 1 Introduction

Clinically, an accurate assessment of drug response in cancer patients is crucial for treatment. Research has demonstrated that changes in genomic profiles significantly impact the effectiveness of cancer therapies and can serve as biomarkers to identify treatment-sensitive populations[1, 2].

Compared to clinical patients who only record the response to one or a few drugs, cell lines cultured in vitro, such as Genomics of Drug Sensitivity in Cancer (GDSC) [1], Cancer Cell Line Encyclopedia (CCLE) [3], provide a large scale dataset for studying the association between drug sensitivity and genome. These datasets are commonly used for pre-clinical modeling, and can also be transferred to patients[4].They offer researchers multi-modal genomic profiles, including mutation (MUT), copy number alteration (CNA), DNA methylation (METH) and gene expression (GE), along with response data to a large number of anticancer drugs.

Quantifying the relationship between tumor genome and drug sensitivity through response scores has become an important and challenging task in bioinformatics and translational medicine[5]. In silico studies providing accurate response predictions are expected to explain resistance mechanisms and guide clinical treatment[6]. Among them, multi-modal fusion methods have been demonstrated to provide comprehensive knowledge of genome and biological processes, therefore improving prediction performance[7–9].

However, complete multi-modal genomic data is not always accessible, and gene expression data is commonly used for analysis in real clinical scenarios. Since multimodal genomic data can be collected from public datasets, the focus of this study is on scenarios where comprehensive genomic knowledge is accessible during the training stage, and only gene expression data is utilized for inference. Among the solutions to missing modalities, knowledge distillation (KD) has emerged as a promising approach to effectively enhance the performance of a single-modality model by transferring knowledge from a multi-modal teacher network[10, 11]. Nevertheless, there are several underexplored challenges in cross-modal genomic KD. Firstly, the quality of the teacher model has a significant impact on the student model, so it is crucial to train a powerful teacher that can not only capture information from each genomic modality but also fully explore genomic relationships. Secondly, in addition to addressing the teacher-student discrepancy on an individual basis during knowledge transfer, it is important to simultaneously transfer inter-sample correlations of genomic features[12, 13].

To address the critical issues mentioned above, we propose a three-stage privileged information distillation framework based on relational perception and feature reweighting. Our aim is to enhance the performance of gene expression data in predicting drug response by leveraging privileged multi-modal genomic knowledge during training. In particular, our work focuses on the above two key problems about KD. First, we integrate the knowledge of multi-modal genomes to improve the existing late-fusion architecture, which not only avoids fusion disasters caused by conflicting modality information, but also establishes the information flow of feature interactions between modalities. Second, we address the issue of losing inter-sample relationship information during the process of KD. We propose a method called relation-aware differential distillation, which aims to efficiently transfer the knowledge of the inter-sample relationships to the student network and consequently enhance the performance of the student model. Experimental results demonstrate that our approach outperforms state-of-theart multi-modal fusion methods using only unimodal gene expression information. Our main contributions are as follows:

- We propose a novel privileged information knowledge distillation framework to learn multi-modal genomic information during training and leverage unimodal gene expression data for prediction.
- To obtain a comprehensive teacher model, we employ a feature fusion attention strategy that benefits from multi-modal genomic information and re-weights feature representations.
- To preserve the inter-sample correlations during distillation and focus on hard samples, we design a relation-aware differential distillation method.

## 2 Related Work

### 2.1 Drug response prediction

Drug response prediction is commonly approached as either a regression or classification problem. In regression analysis, the drug response is typically quantified as the concentration of the drug that reduces cell viability by 50% (IC50). On the other hand, in classification models, continuous IC50 values are discretized into distinct categories, potentially leading to loss of detailed information[6].

In addressing this issue, nonlinear modeling approaches have shown significant advantages in the 2014 Drug Sensitivity Prediction Challenge[14]. Deep learning models have gained popularity due to their robust nonlinear modeling capabilities and have been extensively utilized in recent years for predicting drug responses in cancer cell lines. Without taking into account the impact of prior knowledge on the model, end-to-end deep learning models have been able to generalize to clinical settings[4, 15]. Among them, multi-modal deep learning models such as tCNN[16] and GraphDRP[17] integrate drug information with genomic data to achieve multi-drug learning, enabling simultaneous prediction of drug sensitivity across different drugcell lines. GraOmicDRP[18] and GraTransDRP[19] further incorporate multi-modal genomics data to enhance accuracy. For drug representations, SMILES is commonly used to obtain fingerprints, descriptors, and graph structures. Genomic information typically encompasses MUT, CAN, METH, and GE.

### 2.2 Knowledge Distillation

Knowledge distillation was initially proposed for model compression[20]. By minimizing the KL divergence between the predictions of the large teacher model and the lightweight student model, knowledge can be transferred to the student model to improve its performance. The technical challenges of knowledge distillation can be divided into two aspects: improving the learning ability and transferability of the teacher model. The former focuses on obtaining a more reliable and robust teacher model[21, 22], while the latter aims to achieve better alignment between the teacher model and the student model [23].

## 3 Methods

### 3.1 Data Collection and Preparation

The total number of cell lines or patients included in the dataset, overall counts of drugs, recorded instances of drug response, and data sources are presented in Table 1.

**Table 1.**
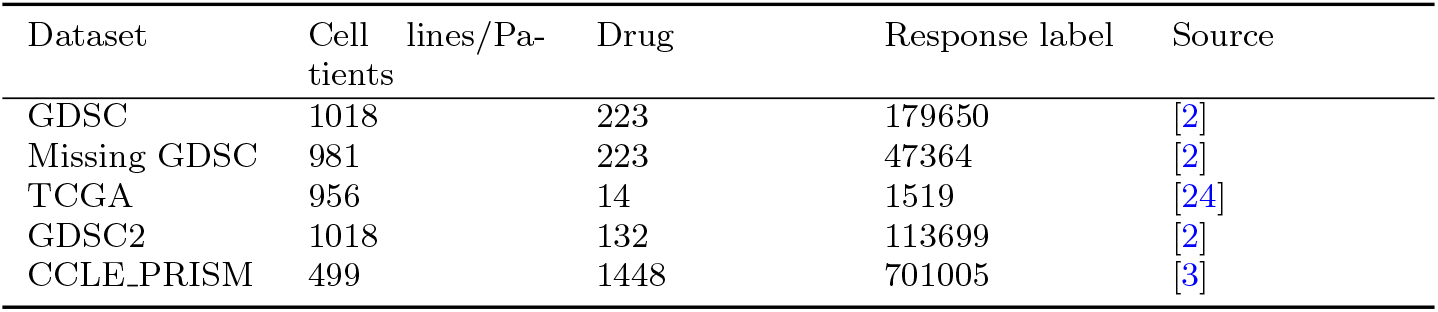
Details of the dataset, including type of cell line or number of patients, type of drug, drug response label, and data source. GDSC dataset is used for training, the other datasets are used for testing or evaluation.

| Dataset | Cell lines/Patients | Drug | Response label | Source |
| --- | --- | --- | --- | --- |
| GDSC | 1018 | 223 | 179650 | [2] |
| Missing GDSC | 981 | 223 | 47364 | [2] |
| TCGA | 956 | 14 | 1519 | [24] |
| GDSC2 | 1018 | 132 | 113699 | [2] |
| CCLE_PRISM | 499 | 1448 | 701005 | [3] |

#### 3.1.1 GDSC Dataset

We collected pharmacogenomic datasets from Genomics of Drug Sensitivity in Cancer (GDSC)1000 project[2], which provides a large number of high-throughput drug screening data across various cancer tissue types. The continuous half-maximal inhibitory concentration (IC50) values were used as metrics for drug sensitivity. Following data preprocessing methods employed in previous studies such as tCNN[16], SCAD[25], GraphDRP[17] and GraTransDRP[19], we focused on 223 drugs tested on 1018 cell lines with drug response IC50 normalized within a range of (0, 1). All genomic data can be downloaded from ^1^.

#### 3.1.2 Missing Dataset of GDSC

We discovered that there are 1018 cell lines and 223 drugs in GDSC, resulting in a total of 227,014 (1018*223) possible combinations. However, GDSC only contains data for 179650 cancer drug responses, leaving 47364 (approximately 20%) pairs missing. As a result, we use our model to predict the unknown cancer drug response.

#### 3.1.3 TCGA Clinical Dataset

In addition, we collected real-world clinical data to assess the effectiveness of our approach on external datasets. The Cancer Genome Atlas (TCGA)[24] provides molecular data along with patients’ clinical information, such as cancer types and drug treatment. It includes data from various cancer types, patient demographic, genomic features, and technical variation. We obtained drug response and gene expression molecular data from here ^2^. After filtering out drugs that were not included in the list of 223 drugs, we only considered data from the remaining patients. This resulted in a total of 956 drug response-gene expression data pairs, consisting of 408 non-responders and 548 responders across 14 drugs.

#### 3.1.4 GDSC2 Dataset

Additionally, We collected 132 unseen drugs and their corresponding drug-genome response dataset (113699 records) from GDSC2. It shares the same genomic profiles with the GDSC dataset, but it validates on new drugs.

#### 3.1.5 CCLE Dataset

We collected the genomic profiles from the CCLE[3], which were then integrated with drug response data from the PRISM Repurposing dataset. This dataset comprises 1,448 compounds screened against 499 cell lines[26]. Notably, it contains a significant number of compounds and genome information that have not been previously observed in GDSC. This dataset is public avaliable on Broad DepMap Portal ^3^.

### 3.2 Problem Settings

We formulate the task of drug response prediction as a regression problem based on neural network. In the teacher model, drug information and genome information are input separately to predict drug response scores for specific interactions between drugs and cell lines. In the student model, only gene expression data and drug information is used for training and inference.

### 3.3 Genome Representations

The gene expression data, denoted as *G*, is stored in matrix form and reduced to 1000 dimensions by Kernel PCA[27] after undergoing *Min-Max Normalization*. Kernel PCA uses kernel functions to map the data to a hidden space, which effectively handles nonlinear mappings and prevents the interference of redundant information for network convergence. DNA changes at a single base or fragment, as well as copy number variation, are uniformly classified as gene mutations. DNA sequence changes are denoted as 1, while no changes are denoted as 0, resulting in the gene mutation matrix *M*. The DNA methylation data is also represented by a binary matrix *E*, indicating whether the gene is hypermethylated (1) or hypomethylated (0). Additionally, cell lines with missing modalities are removed and the remaining data is used for training and validation. We implement an end-to-end convolutional neural network (CNN)[19] for gene feature extraction without manual intervention. Finally, the extracted feature maps of multi-channel hidden space are denoted as *f* (*G*), *f* (*M*), *f* (*E*) respectively.

### 3.4 Drug Molecular Representations

A graph-transformer network is designed to learn the representations of molecular chemical structure[19]. The input comprises graph features represent as *G*_*d*_ = (*N*_*d*_, *E*_*d*_), where 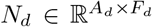 denotes the node matrix of the graph, corresponding to atoms within the molecular structure. 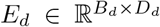 represents the adjacency matrix of edges, describing the connections between the nodes, i.e., the chemical bonds. Where *A*_*d*_ denotes the number of atoms in the molecule, *B*_*d*_ represents the number of chemical bonds connecting these atoms. Additionally, *F*_*d*_ refers to the feature embedding of atomic properties while *D*_*d*_ refers to bond endpoints and is consistently set to 2 due to each bond connecting two atoms. This combined architecture of a Transformer with a graph neural network effectively represents the chemical structure of the drug. This approach leverages Transformer’s capability in handling long-range dependencies within sequential SMILES data[28]. Simultaneously, the Graph Neural Network[29] component reveals the intricate inter-connectivity among atoms, thus reflecting actual spatial structures and bonding relationships in molecular.

### 3.5 Framework

#### 3.5.1 Privileged Information KD Framework

In the context of cross-modal KD, information tends to transfer from a well-annotated modality to another where labels may not always be available. Inspired by privileged information learning[30, 31], we modify the cross-modal distillation framework into a privileged information KD framework to completely exploit the potential of the available data during training. Our approach involves training the network through a three-step learning process (Figure 1). In the initial stage, we employ a late-fusion multi-modal network[17] as the pre-trained model for the teacher and an unimodal network[17] for the student. Both models consist of two components: a feature extractor and a feature regressor. In the second stage, feature map 1 is obtained by integrating the hidden representations of gene expression and drug streams, while feature map 2 is obtained by integrating the hidden representations of other genomic data and drug streams. These two feature maps are then input into different regressor for regression scoring. In contrast to late fusion, our approach separates gene expression information from other genomic data during training in order to avoid spurious learning caused by conflicting information between different modalities. Concurrently, as different modalities often have non-equilibrium effects on the final result[32], this approach prevents any single modality from dominating and ensures that information from other modalities is not overshadowed during training. The two regression scores *y*_1_, *y*_2_ are combined with equal weights to calculate the final prediction *ŷ*. The Mean Squared Error (MSE) loss is defined as the discrepancy between the predicted value and the actual label *y*:

**Fig 1.**
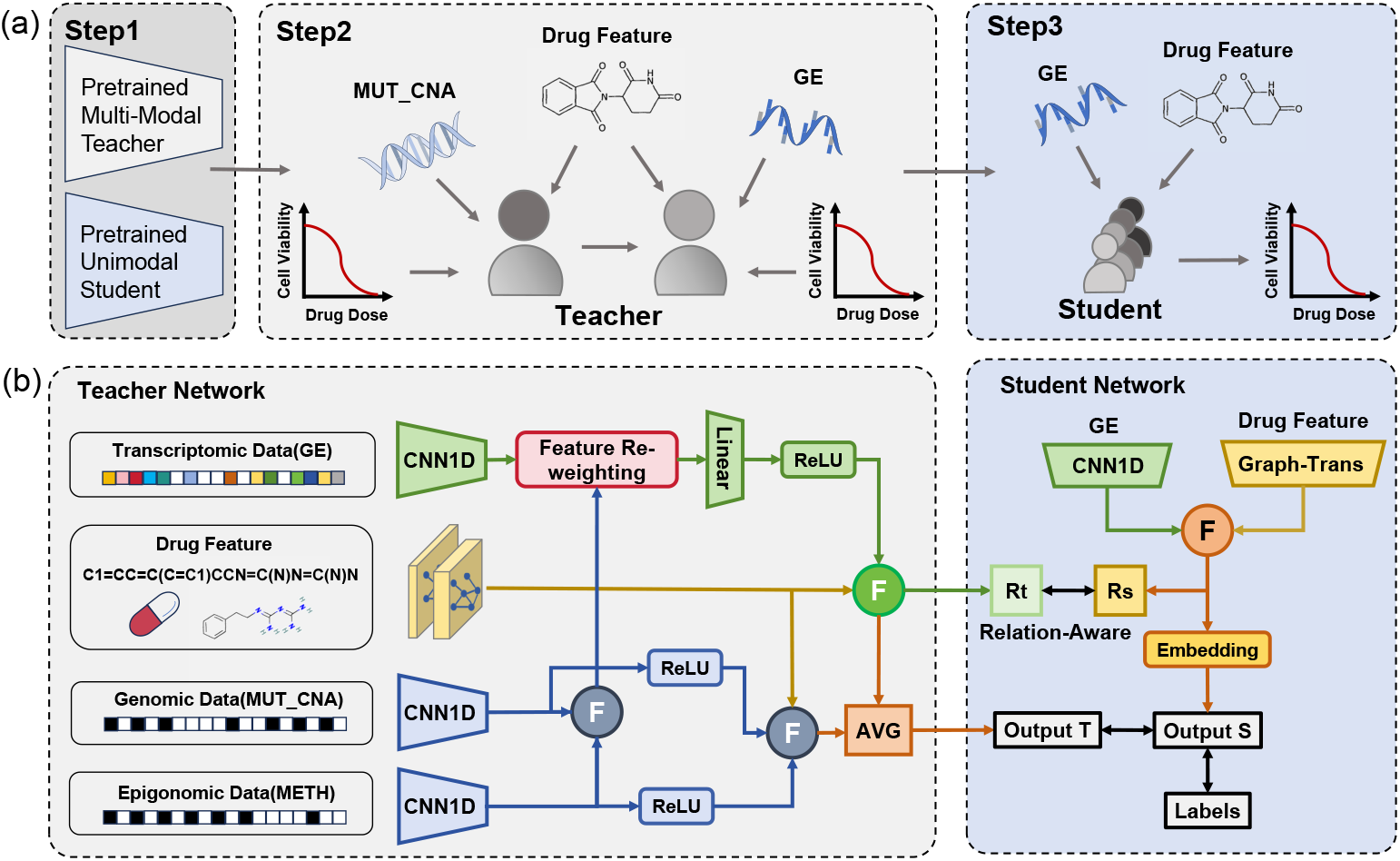
An overview of our proposed privileged KD framework. (a)Three steps of distillation: (i) Pretrain a multi-modal teacher and an unimodal student network. (ii) Train a feature re-weighting multi-modal teacher network. (iii) Distill the multi-modal teacher network to the student by KD loss or RAD loss. (b) The structure of our framework includes the teacher with multi-modal genomic data as inputs and the student with only gene expression data as input. MUT: mutation, CNA: copy number alteration, METH: DNA methylation and GE: gene expression.

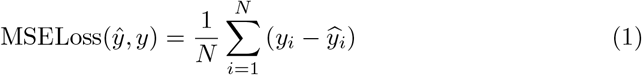

In the second stage, we use the parameters of the feature extraction in the first stage to initialize the parameters of the network to accelerate convergence.

In the third stage, the prediction *y*_*t*_ of the teacher network is aligned with the output *y*_*s*_ of the student network. In classification models, student predictions are typically guided by soft labels of teachers obtained by setting temperature parameters. This label smoothing may provide additional information beyond just the probability distribution over individual classes. However, in regression tasks, the output labels consist of inherently continuous values. Therefore, we directly process the soft labels as the output of the teacher network and apply *MSE* loss for alignment[33]. This is also denoted as the *KD* loss:

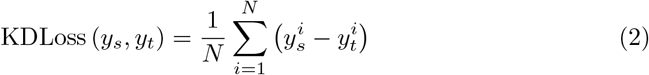

In addition, overfitting the teacher’s knowledge without differentiation may mislead students. It is necessary to regularize the student network and introduce a label regularized *MSE* loss:

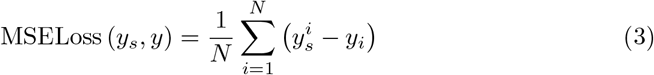

This regularization loss aims to reduce the complexity of the model and effectively mitigate the risk of overfitting for both simple and difficult samples. Moreover, we have also adopted *Dropout* and *L2* regularization to further reduce network complexity.

#### 3.5.2 Feature Reweighting for Teacher Learning

In order to obtain a robust teacher model, it is essential to consider inter-modality correlations in addition to mining information from each modality. The associations between modalities can provide additional knowledge injection, establishing a close relationship between the two relatively independent streams established in Section 3.5.1. Since gene expression information is utilized for final inference, we have chosen to incorporate mutation and methylation knowledge into the gene expression data stream. After extracting deep features from raw genome data by convolutional kernels, feature maps with the same number of channels but different feature dimensions are obtained as 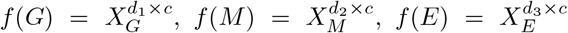. The methylation profile is concatenated with the gene mutation profile to obtain a new profile 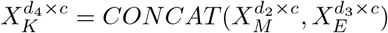 (genomic profiles).

Inspired by the cross-attention mechanism between different sequences in NLP tasks[34, 35], we calculate the similarity between genomic features and transcriptome features. We then re-weight the transcriptome features to ensure that the final output feature map contains complementary information from both modalities. Particularly, the feature maps from genomic profiles are transformed by the projection matrix 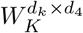 and 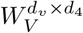, to obtain 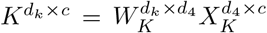 as key and 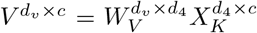 as value. While the feature map obtained from gene expression is transformed by the projection matrix 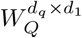 to obtain the feature map 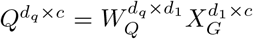, which is regarded as the query set. After the cross-attention process, each element of the attention map represents the feature correlation between the query set and the key set. This is accomplished by initially computing the dot product between the query vectors and the key vectors and outputting the attention map. Subsequently, this matrix is typically normalized by dividing it by the square root of the dimension of the key vectors (often represented as 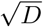) to ensure that the values remain within a reasonable range, thereby preventing issues with gradients during training. Then, the original feature map 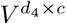 is re-weighted using the attention graph, allowing each channel of the output feature map to contain information from all other channels. In particular, channels corresponding to features with strong correlations are assigned relatively large weights. To mitigate potential biases and ensure robustness, the cross-attention mechanism dynamically calibrates the feature weights, preventing overemphasis on any modality. Finally, the average of the prediction of gene expression stream and other genome streams is calculated as the final output. This model is referred to as averaged net (*AVG NET*). As a result of cross-attention, the feature map 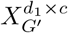 has the same dimension as the gene expression feature map 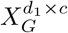, with each feature dimension being re-weighted accordingly:

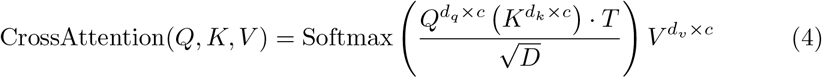

where *D* denotes the dimension of hidden layer, which is typically equal to *d*_*k*_.

#### 3.5.3 Relation-Aware Differential Distillation

The distillation method mentioned in *Section 3.5.1* aligns the predictions of the student and teacher models equally across samples, disregarding inter-sample correlations and differentiation. In this part, we will introduce the detail implemention of relation-aware differential distillation. First we discuss the alignment of inter-sample correlations. Due to the rich information contained in the hidden space after feature extraction, we consider the correlation of the hidden space features as the intersample correlation. The gene expression hidden space 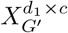 obtained after feature reweighting in *Section 3.5.2* was used as the meta knowledge, which contains valuable genomic information. Here, we utilized meta-knowledge as feature vectors to calculate the inter-sample correlation. Due to its computational efficiency and compatibility with softmax, the dot product method was employed to measure the inter-sample correlation. The *MSE* loss was applied to quantify the dissimilarity of this correlation between the student and the teacher so that knowledge difference can be narrowed by distillation:

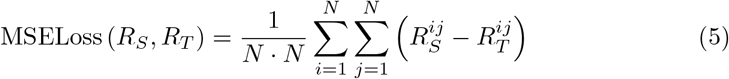

where 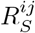 denotes the correlation matrix of feature vectors output by the student network, 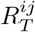 denotes the correlation matrix of feature vectors output by the teacher network.

Secondly, we need to consider the heterogeneity of the samples. What students learn from their teachers actually represents a knowledge gap, and this gap often varies among different samples[36]. The problem of hard sample mining in computer vision is indeed a challenging task. These samples with higher loss values generally have a significant impact on the final performance. Drawing inspiration from this concept, we define samples with a large knowledge gap between the student and the teacher as hard samples, which typically require more time for the student to learn. This knowledge gap can be simply expressed as *O*_*T*_ − *O*_*S*_. Where *O*_*T*_ denotes as the output of the teacher and *O*_*S*_ denotes as the output of the student. The overall structure of our relation-aware differential distillation module is illustrated in Figure 2.

**Fig 2.**
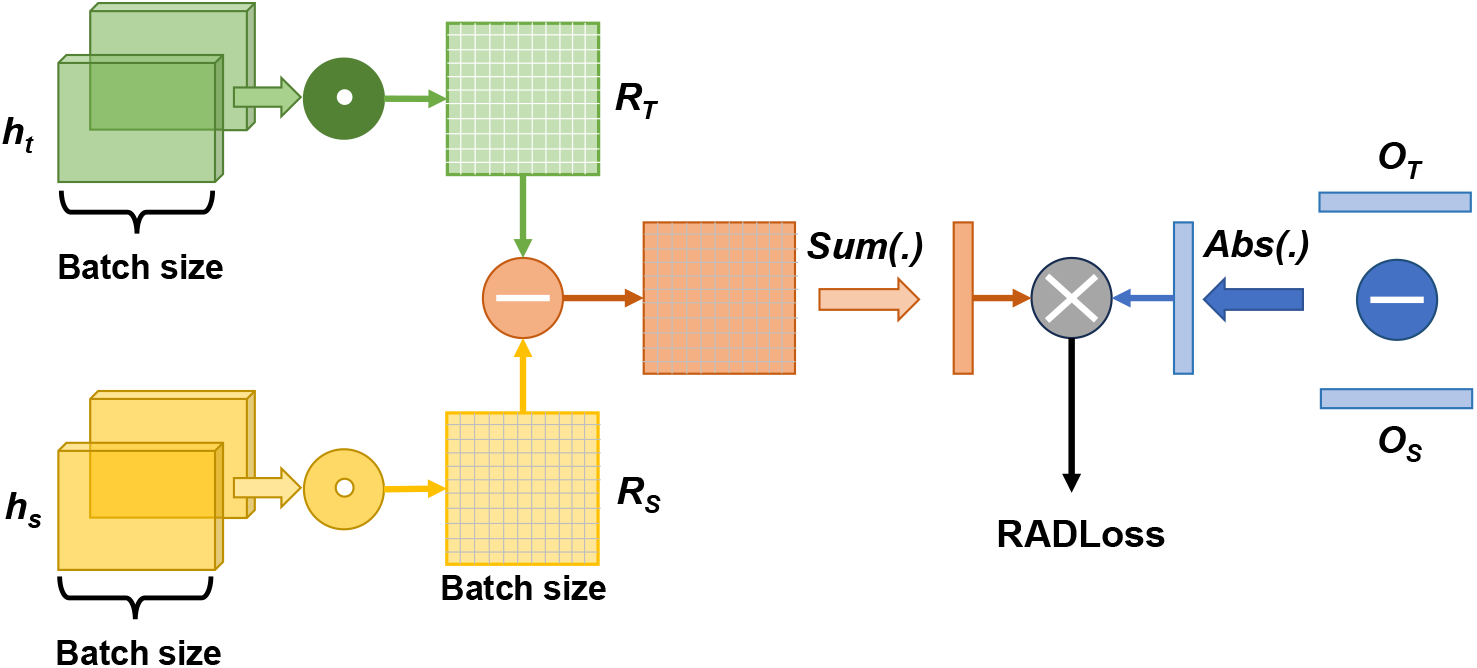
Relation-aware differential distillation module. *R*_*S*_ and *R*_*T*_ denote the relation-aware matrices. *O*_*T*_ and *O*_*S*_ represent the output of the teacher and the student, respectively.

Finally, we multiply the aforementioned two components to obtain the loss for relation-aware differentiation (RAD) distillation:

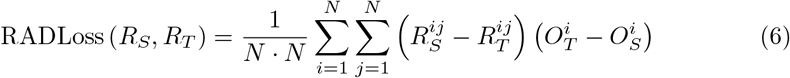

This means for hard samples, it is essential to prioritize the acquisition of relational knowledge and therefore assign greater weights. Conversely, for simple samples, acquiring relational knowledge is relatively effortless, resulting in smaller weights. As *R*_*S*_, *R*_*T*_, *O*_*S*_, and *O*_*T*_ are obtained through non-parametric computations, the *RADLoss* function does not directly involve any additional model parameters.

Similarly, the regularization loss mentioned in *equation 3* is also added to avoid overfitting.

## 4 Results

### 4.1 Model Evaluation

#### 4.1.1 Evaluation Metric

We employed root mean squared error (RMSE) and Pearson correlation coefficient (PCC) to assess the performance of the regression model. *RMSE* measures the discrepancy between the actual and predicted values, while *PCC* is a widely used metric for evaluating correlation. The formulas for these indicators are as follows:

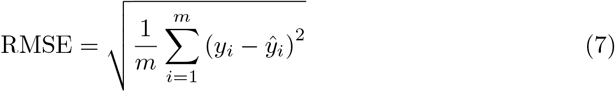

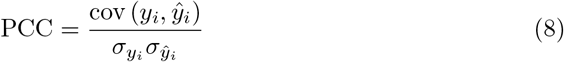

where *y*_*i*_ and *ŷ*_*i*_ represent the true value of IC50 and the predicted IC50, respectively, with *σ* represents the standard deviation.

In the ablation experiment, we also calculated the raw RMSE and raw PCC, which are the restoration of the predicted normalized IC50 value to its original IC50 value. The logical function for normalizing the IC50 (*x*) is expressed as:

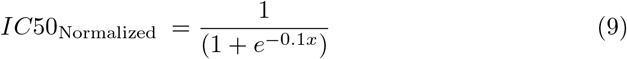

where the inverse transformation represents the IC50 value as labeled in the dataset.

#### 4.1.2 Baselines

We found four representative baselines with the same data source, all of which were not trained with additional knowledge. They integrated genomic data with drug information to model the nonlinear relationship between fused information and drug response. Among them, tCNN[16] and GraphDRP[17] use gene mutation data, while GraomicDRP[18] and GraTransDRP[19] incorporated all modalities. For each set of experiments, the results from three repeated randomized trials were averaged, all exhibiting variances below 10-5; therefore, the display of variance was omitted.

### 4.2 Implementation details

Our approach is implemented with the PyTorch library[37]. We employ the same feature extractor backbone as in previous work[19]. The drug stream consists of two transformer layers with two graph layers: a GCN layer and a GAT layer. Based on prior research, the genome stream consists of three one-dimensional convolutional neural networks (1D CNNs) with a kernel size of 8 and channel dimensions of 32, 64, and 128. The concatenated layer is a fully connected layer with input channels determined by the input modality and output channels set to 1024. The regressor consists of two fully connected layers, one with dimension 1024 and the other with dimension 128. We employ a grid search strategy to identify the optimal learning rate for network optimization. The learning rate is varied within the range of 1e-3 to 1e-5. Additionally, the training process is constrained to fewer than 300 epochs through an early stopping strategy with a patience parameter set to 5. We utilize the Adam optimizer with *β*_1_ =0.9 and *β*_2_ =0.99 for optimization purposes. The batch size is determined by the dataset and the device, which we set to 512 in our experiments.

### 4.3 Experimental Results

As depicted in Table 2, our findings on the test set are as follows: (1) Compared with all single-modal gene expression predictions, our student network improved by approximately 6% over the best baseline, reaching RMSE of 0.0234 and PCC of 0.9372. (2) In comparison to all multi-modal models, our multi-modal teacher network exhibits a 3% improvement over the baseline model, with an RMSE of 0.0232 and a PCC of 0.9378, outperforming the distilled single-modal network. It shows that reweighting feature representation strategies that benefit from multi-modal genomic information can yield more comprehensive teacher models. It can completely mine the information of each stream, surpassing late fusion-based multi-modal approaches. (3) Notably, our student networks outperformed the best multi-modal baseline networks, suggesting that making predictions using just one modality is feasible. These results imply that better teachers tend to produce better students; although the superior unimodal student does not outperform the superior multi-modal teacher, it does outperform the general teacher.

**Table 2.**
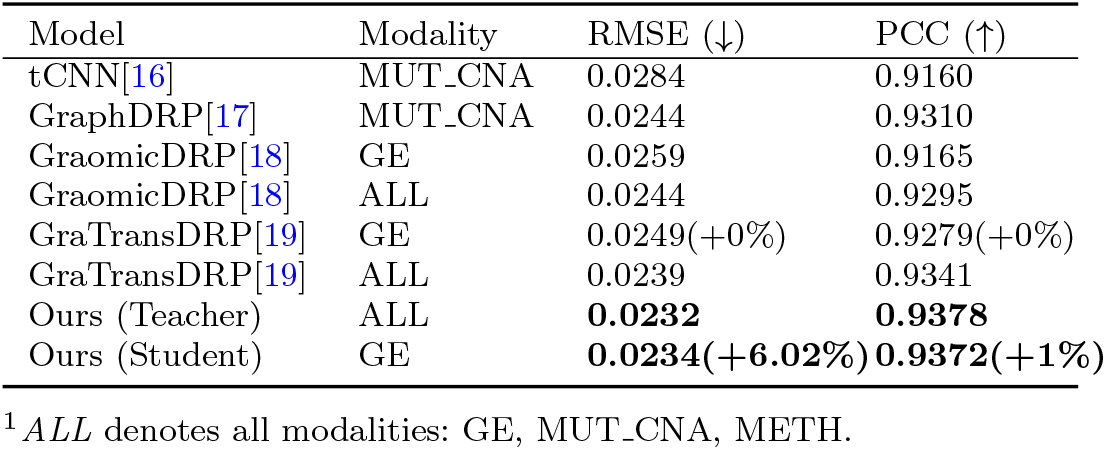
Comparison of feature re-weighted teacher model and distilled student model with sota baselines on GDSC dataset. The proposed teacher model performs best, and the student network outperforms the multimodal networks of the other methods.

### 4.4 Ablation Study

#### 4.4.1 Evaluate the Contributions of Each Strategy

We perform an ablation experiment to evaluate the individual contributions of the strategies. As shown in Table 3, the first row adopts the existing multi-modal fusion method[19]. In the second row, we apply KD between the multi-modal network obtained in Experiment 1 and the single-modal network. We observe that the distilled single-modal model (row2) outperforms the late-fusion multi-modal network (row1). Late-fusion multi-modal networks may experience information conflicts between different modalities due to data noise, leading to a reduced exploitation of modality information. Conversely, in a single-modal student network, there are no such information conflicts between modalities. In contrast, in row 3, we employ a multi-modal teacher trained with our proposed privileged information KD framework. The superiority of row 3 over row 1 demonstrates that this framework allows for more comprehensive exploration of multi-modal knowledge. The fifth row represents the direct distillation from the teacher network obtained in Experiment 3 to the unimodal student network. The superiority of row5 over row2 illustrates that our proposed multimodal teacher network can provide better guidance for student learning. As shown in row 4, applying our feature re-weighting strategy further improves the performance of our proposed multi-modal teacher. Distilling from this synthetic learning teacher, the student module (row5) improves RMSE by 0.86% over the vanilla GE module (row2). Compared with row6 and row7, row8 demonstrates the advantage of relation-aware differential distillation in distilling inter-sample correlations from teachers and learning knowledge gaps heterogeneously. In order to intuitively reflect the effectiveness of students learning the inter-sample relationship from the teacher model, we calculate the feature similarity among each batch of samples (batch size=256). The correlation matrix in Figure 3(b) reflects the distance between the teacher and student. It is evident that our RAD distillation method results in a smaller teacher-student gap compared to the baseline (W/O KD) and KD only method, indicating that our approach distills the richest inter-sample knowledge from the multi-modal teacher. The linear scatter plots of our final unimodal drug response prediction are presented in Figure 3(a). A closer scatter to the diagonal line indicates a closer model prediction to the true value.

**Table 3.**
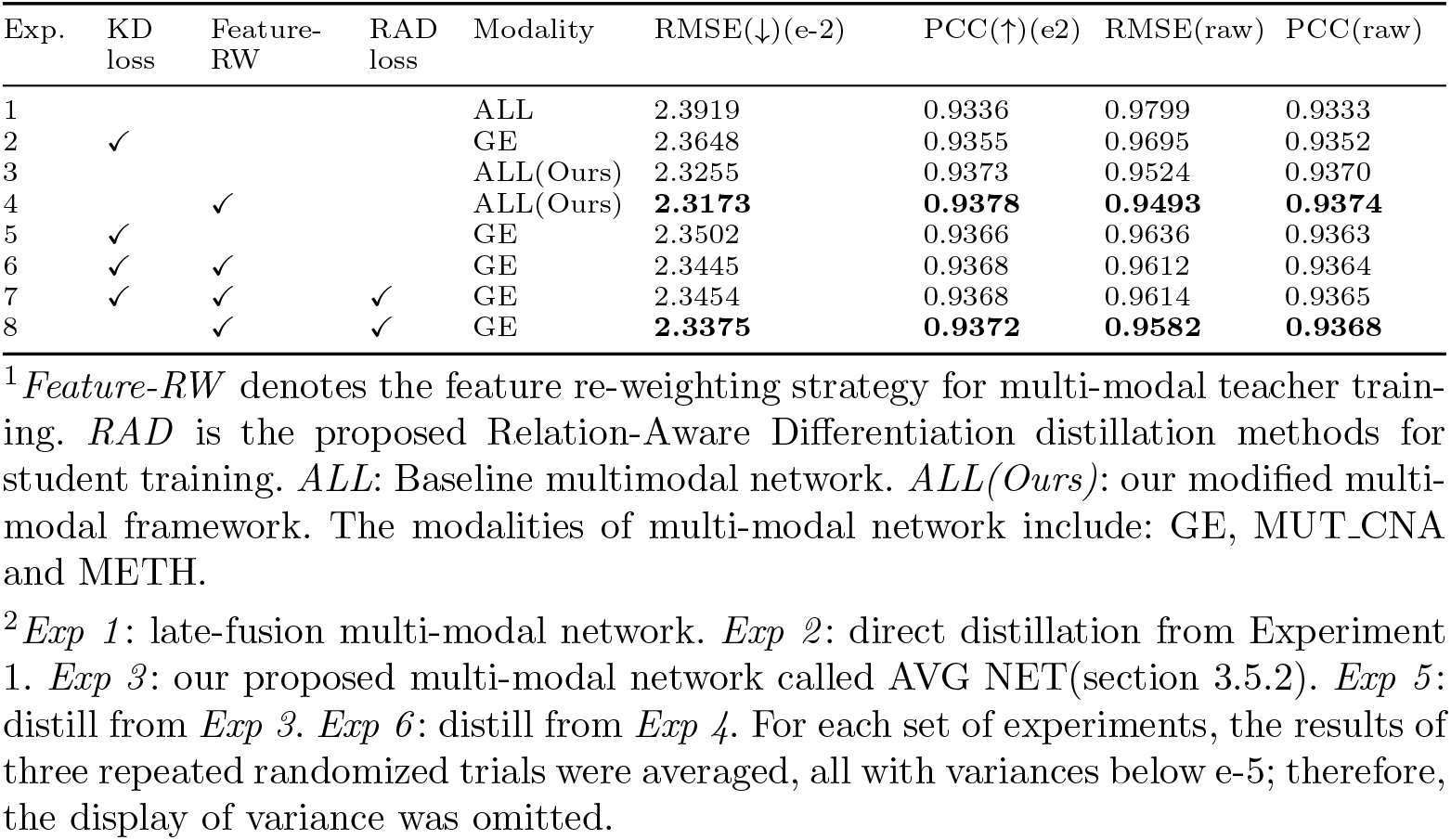
Ablation study demonstrated that the combination of feature re-weighting and relation-aware differentiation distillation results in the best-performing student on the test set.

| Exp. | KD loss | Feature-RW | RAD loss | Modality | RMSE(↓)(e-2) | PCC(↑)(e2) | RMSE(raw) | PCC(raw) |
| --- | --- | --- | --- | --- | --- | --- | --- | --- |
| 1 |  |  |  | ALL | 2.3919 | 0.9336 | 0.9799 | 0.9333 |
| 2 | ✓ |  |  | GE | 2.3648 | 0.9355 | 0.9695 | 0.9352 |
| 3 |  |  |  | ALL(Ours) | 2.3255 | 0.9373 | 0.9524 | 0.9370 |
| 4 |  | ✓ |  | ALL(Ours) | **2.3173** | **0.9378** | **0.9493** | **0.9374** |
| 5 | ✓ |  |  | GE | 2.3502 | 0.9366 | 0.9636 | 0.9363 |
| 6 | ✓ | ✓ |  | GE | 2.3445 | 0.9368 | 0.9612 | 0.9364 |
| 7 | ✓ | ✓ | ✓ | GE | 2.3454 | 0.9368 | 0.9614 | 0.9365 |
| 8 |  | ✓ | ✓ | GE | **2.3375** | **0.9372** | **0.9582** | **0.9368** |

<sup>1</sup>*Feature-RW* denotes the feature re-weighting strategy for multi-modal teacher training. *RAD* is the proposed Relation-Aware Differentiation distillation methods for student training. *ALL*: Baseline multimodal network. *ALL(Ours)*: our modified multi-modal framework. The modalities of multi-modal network include: GE, MUT.CNA and METH.
<sup>2</sup>*Exp 1*: late-fusion multi-modal network. *Exp 2*: direct distillation from Experiment 1. *Exp 3*: our proposed multi-modal network called AVG NET(section 3.5.2). *Exp 5*: distill from *Exp 3*. *Exp 6*: distill from *Exp 4*. For each set of experiments, the results of three repeated randomized trials were averaged, all with variances below e-5; therefore, the display of variance was omitted.

**Fig 3.**
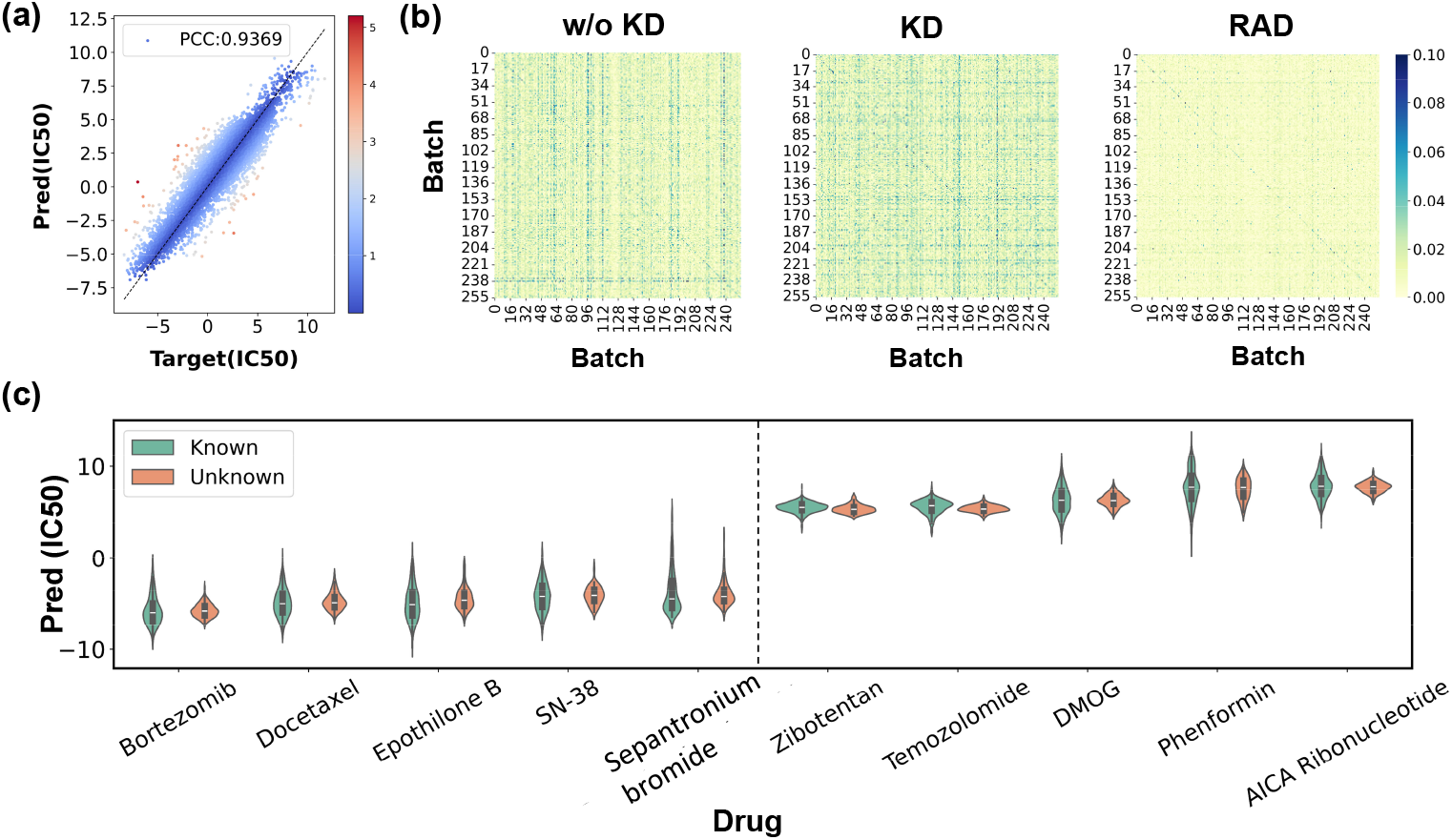
Visualization of drug response prediction performance on the GDSC and missing data sets. (a) The linear scatter plot illustrates our final unimodal drug response prediction results, with color indicating the distance between the predicted IC50 and the true value. (b) Visualizing the distance between the correlation matrix of the teacher network and the student network. “W/o KD” refers to the original pre-trained multi-modal teacher and unimodal student, while “KD” refers to applying MSE distillation loss between teacher and student. “RAD” refers to implementing our proposed relation-aware differential distillation loss. It demonstrates that our module facilitates more effective knowledge transfer and enhances the alignment of inter-sample correlation. The correlation matrix represents the correlation between features of a batch of samples (batch size = 256). (c) The IC50 distributions for the top-5 sensitive and top-5 resistant drugs in the missing data set are consistent with the GDSC dataset. ‘Known’ indicates labeled GDSC dataset of each drug, while ‘Unknown’ indicates missing labeled data of each drug. This confirms our model’s ability to discriminate between response concentrations of different drugs across cell lines.

#### 4.4.2 Sensitivity Analysis Experiments on Feature Re-weighting

Methods that compute feature re-weighting may yield three distinct weighting outcomes, utilizing gene mutation, gene expression, and methylation features as query objects, respectively. Our experimental findings demonstrate that employing gene expression features as query objects results in the most effective feature re-weighted teacher model on the test set. The experimental results are presented in Table 4.

**Table 4.** The sensitivity analysis experiment of query features. Employing gene expression features as query objects results in the most effective feature re-weighted teacher model on the test set.

| Query | RMSE(e-2)(↓) | PCC (↑) | RMSE(raw) | PCC(raw) |
| --- | --- | --- | --- | --- |
| GE | **2.3173** | **0.9378** | **0.9493** | **0.9374** |
| MUT | 2.3585 | 0.9355 | 0.9659 | 0.9352 |
| METH | 2.3556 | 0.9356 | 0.9651 | 0.9353 |

We further conducted sensitivity analysis experiments on GE-based cross-modality attention. The similarity function is a crucial parameter of the cross-attention mechanism that impacts the feature re-weighting process. We employed three general methods to compute similarities and assessed their performance on the multimodal teacher network, including dot product, cosine similarity and Euclidean distance. (Table 5). The result shows that dot product perform best on the test set.

**Table 5.** Sensitivity analysis experiments for cross-attention. The results show that the Dot product performs the best on the test set.

| Similarity function | RMSE(e-2)(↓) | PCC (↑) |
| --- | --- | --- |
| Base(W/o feature re-weighting) | 2.3919 | 0.9336 |
| Dot product(Ours) | **2.3173** | **0.9378** |
| Cosine similarity | 2.3658 | 0.9350 |
| Euclidean Distance | 2.3777 | 0.9344 |

### 4.5 Scalability Evaluation

Based on the proposed optimal pre-training framework in this paper, we validate the scalability of the model across various scenarios. The additional settings examined include unimodal gene mutation sequencing, unimodal methylation sequencing, and combined sequencing of any two modalities: *MUT CNA&GE, METH&GE*, and *MUT CNA&METH*. We evaluate their performance improvements following the implementation of the pre-training and distillation framework outlined here. The results are summarized in Table 6. The findings indicate that distillation from the proposed multimodal teacher model enhances performance overall when compared to training from scratch.

**Table 6.** The scalability evaluation experiment. The network demonstrates adaptability to any combination of modalities across various scenarios.

| Model | Test Mods | RMSE(e-2)(↓) | PCC (↑) | RMSE(raw) | PCC(raw) |
| --- | --- | --- | --- | --- | --- |
| Baseline | MUT.CNA | 2.3586 | 0.9356 | 0.9665 | 0.9353 |
| Ours | MUT.CNA | **2.3352** | **0.9375** | **0.9573** | **0.9372** |
| Baseline | METH | 2.6545 | 0.9175 | 1.0868 | 0.9172 |
| Ours | METH | **2.6323** | **0.9191** | **1.0777** | **0.9189** |
| Baseline | MUT.CNA&GE | 2.3866 | 0.9339 | 0.9777 | 0.9336 |
| Ours | MUT.CNA&GE | **2.3543** | **0.9362** | **0.9650** | **0.9359** |
| Baseline | METH&GE | 2.3933 | 0.9335 | 0.9803 | 0.9332 |
| Ours | METH&GE | **2.3549** | **0.9360** | **0.9651** | **0.9357** |
| Baseline | MUT.CNA&METH | 2.3587 | 0.9356 | 0.9665 | 0.9353 |
| Ours | MUT.CNA&METH | **2.3332** | **0.9374** | **0.9565** | **0.9371** |

<sup>1</sup>Baseline: training from the scratch.
<sup>2</sup>Ours: distilling from the proposed multi-modal teacher model.

### 4.6 Transferability on Missing Data

As described in Section 3.1.2, we make predictions for 47364 missing cancer drug response data points within the GDSC dataset. Figure 3(c) displays the top-5 sensitive and top-5 resistant drugs, ranked by their average IC50 values. Additionally, violins are plotted against the labeled data from the original dataset, revealing a similarity in the distribution of missing data to that of labeled data. The data set encompasses a total of 14 tissues and 58 tissue subtypes, indicating that some of the missing cell lines belong to the same tissue as those that are known. While we cannot assess the numerical accuracy of our unlabeled predictions, it is reasonable to expect consistency in drug response among similar tumors when treated with the same drug. This indicates our model’s capability to infer responses for new drug-cell line combinations. Notably, Bortezomib demonstrates a tendency towards sensitivity in cancer cell lines, which aligns with findings from Jiang et al[38]. In addition to its significant efficacy against multiple myeloma and certain lymphomas[39, 40], literature also reports its effectiveness against other solid tumors[41, 42] and hematological tumors[43].

### 4.7 Transferability on Clinical Data

In this study, only gene expression data was preformed to predict drug sensitivity, yielding continuous IC50 values. However, the clinical drug response data provided by TCGA consists of four discrete states: progressive disease (PD), stable disease (SD), partial response (PR) and complete response (CR). In this study, PD and SD patients were treated as non-responders, while PR and CR patients were treated as responders. The IC50 value in this study represents a point on the dose-response curve obtained by fluorescent-based cell viability assay, indicating the drug concentration required to suppress half of the cell lines. A higher IC50 value indicates stronger resistance.

To validate the performance of the distilled student model on external datasets, all parameters are frozen for inference.

In this section, we specifically investigate the model’s capacity to differentiate between response concentrations of various drugs in populations. We screen 12 drugs with more than 15 patients. The average predicted IC50 values are then ranked by drug, with the top 3 ‘sensitive’ and top 3 ‘resistant’ displayed in Figure 4(a) and Figure 4(b). The Mann-Whitney test was conducted to compare the distribution of IC50 between the sensitivity and the resistance, resulting in a total of 9 (3*3) outcomes (Figure 4(c)). The results indicate that all p-values are below 0.0001, suggesting that sensitive drugs may have a lower IC50 based on alternative hypotheses. This result is consistent with that of the GDSC dataset. According to the TCGA data, the overall response rate of Vinblastine (n=16) for patients suffering from Bladder Cancer (BLCA), Lung Adenocarcinoma (LUAD), and Skin Cutaneous Melanoma (SKCM) is 9/16. In contrast, the response rate of Bicalutamide (n=17) for patients suffering from Prostate Adenocarcinoma (PRAD) is only 3/17. The IC50 distribution for the GDSC dataset is also depicted, showing that the response levels of these six drugs appear to be consistent across both datasets. For the GDSC dataset, there is minimal overlap between the IC50 distributions of these two drugs (Figure 4(b)), with Vinblastine having a significantly lower IC50 than Bicalutamide. Validation on TCGA clinical data demonstrates the ability of our model to distinguish response levels to different drugs.

**Fig 4.**
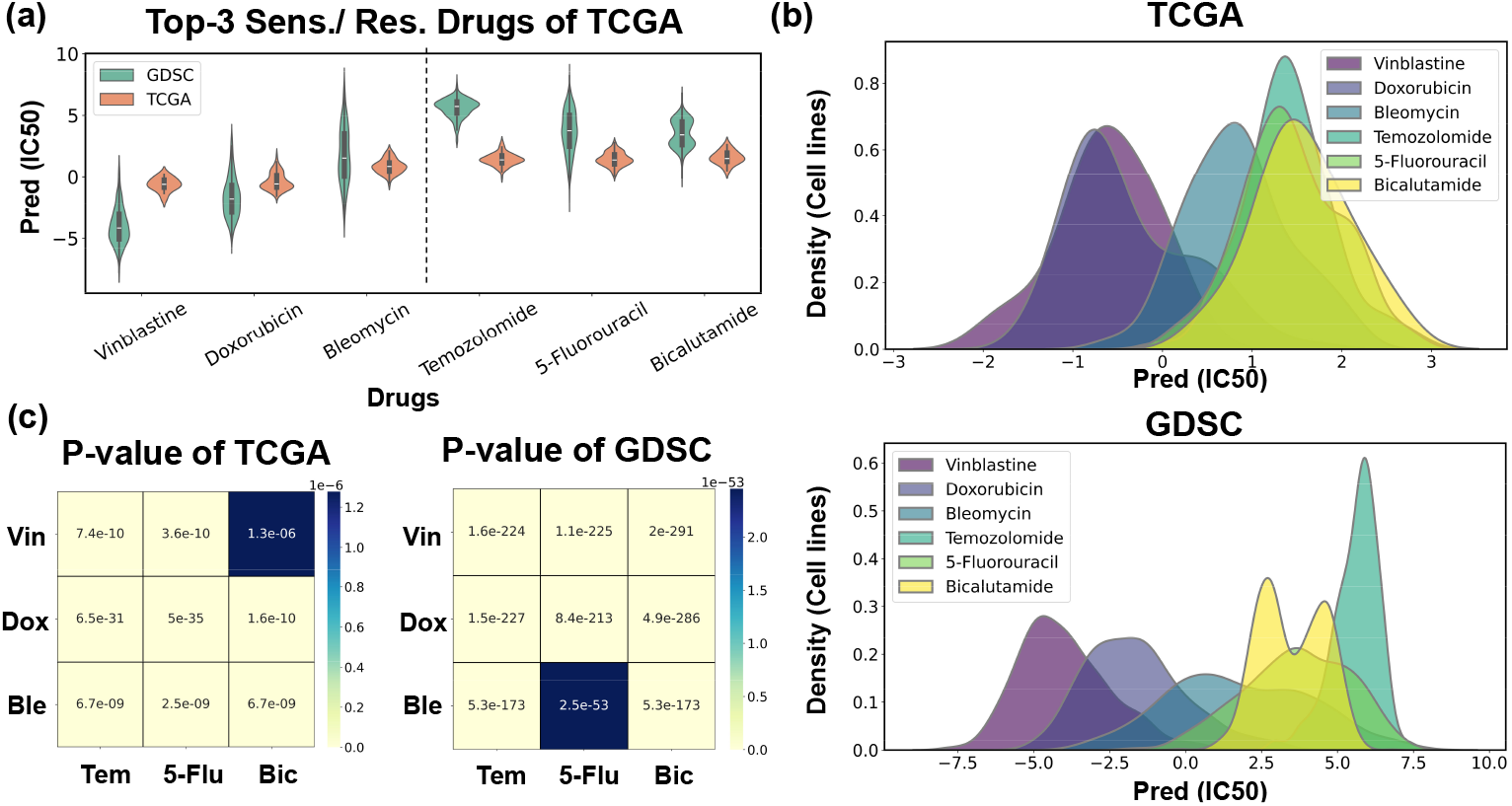
Investigating our model’s capability to differentiate between response concentrations of various drugs in populations. (a) (b) Distribution of IC50 for the top-3 ‘sensitive’ (Vinblascine, Dox-orubicin, Bleomycin) and top-3 ‘resistant’ (Temozolomide, 5-Fluorouracil, Bicalutamide) drugs in TCGA and the drug response distribution for these drugs in GDSC. (c) The Mann–Whitney tests between the ‘sensitive’ drugs and the ‘resistant’ drugs indicate that ‘resistant’ drugs may have a higher IC50 based on the alternative hypothesis (p-value¡0.0001), which is consistent with the GDSC dataset. Vin: Vinblascine, Dox: Doxorubicin, Ble: Bleomycin, Tem: Temozolomide, 5-Flu: 5-Fluorouracil, Bic: Bicalutamide.

### 4.8 Transferability Across Other Cell Line Datasets

We compared our pretrained unimodal network (after privileged information knowledge distillation) with random initialized and original pretrained unimodal network. After fine tuning, we plotted the results of 20 epochs (Figure 5). The results indicate that our model demonstrates rapid convergence and superior generalization capabilities on new data, outperforming random initialization and the baseline. After 10 epochs of fine-tuning, the PCC reached 0.8738 on the GDSC2 test set and 0.7959 on the CCLE test set (Table 7).

**Table 7.** Results are presented on the test sets of GDSC2 and CCLE following fine-tuning for 10 epochs. These findings indicate that transfer learning from our pre-trained model can significantly enhance generalization performance on the test set.

| Dataset | Model | MSE(e-2)(↓) | PCC (↑) |
| --- | --- | --- | --- |
| GDSC2 | Random initialization | 0.1146 | 0.8392 |
| GDSC2 | Base unimodal network | 0.0734 | 0.8710 |
| GDSC2 | Ours | **0.0702** | **0.8738** |
| CCLE | Random initialization | 0.1331 | 0.7953 |
| CCLE | Base unimodal network | 0.1287 | 0.7957 |
| CCLE | Ours | **0.1256** | **0.7959** |

<sup>1</sup> *Ours* denotes our pretrained unimodal network after privileged information knowledge distillation.

**Fig 5.**
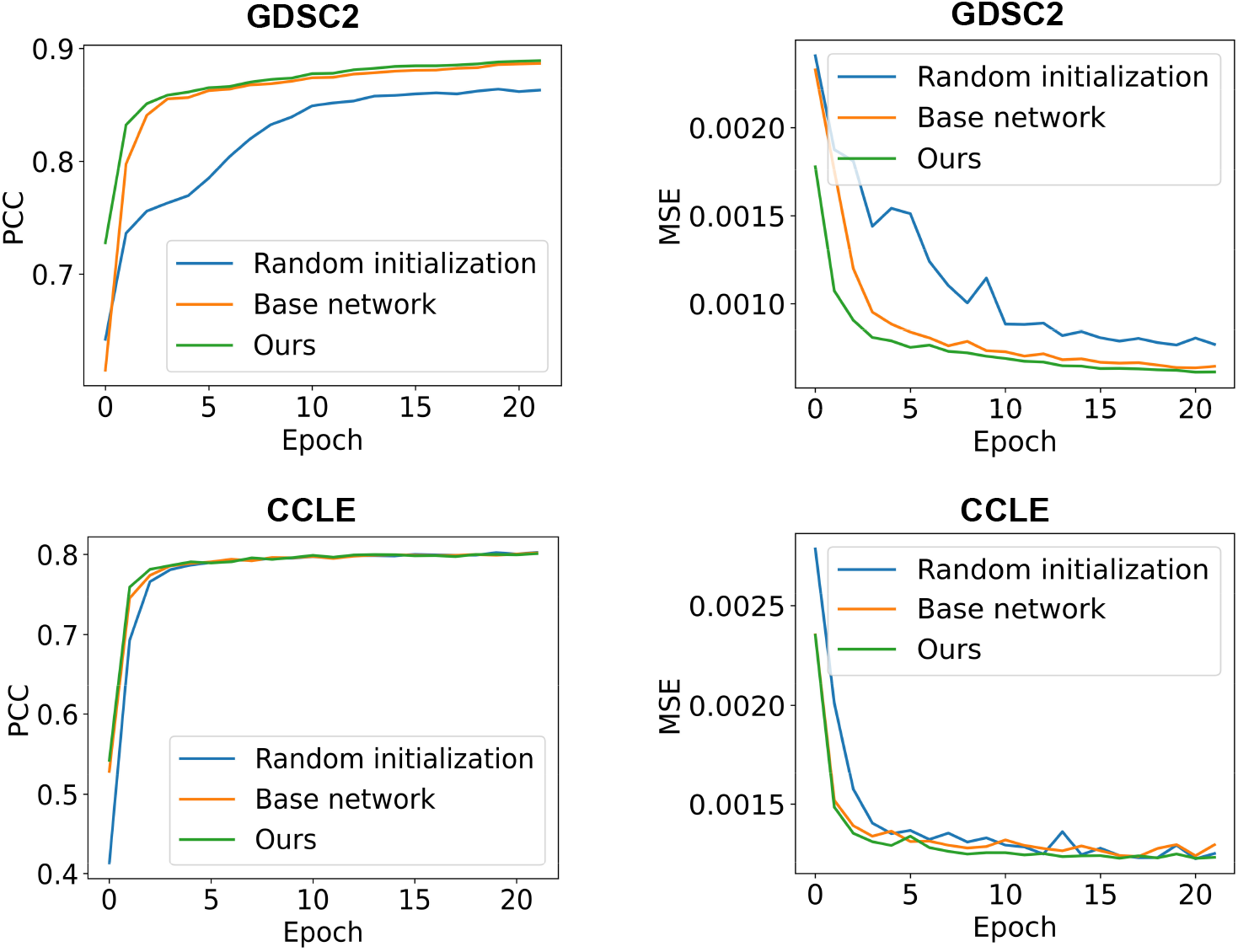
PCC and MSE on the test sets of GDSC2 and CCLE. Our model demonstrates rapid convergence and superior generalization capabilities on new data, outperforming random initialization and the baseline.

## 5 Discussion

The resistance mechanism is highly intricate and involves numerous factors. Despite pre-training based on data from cancer cell lines cultured in vitro, there are still significant challenges for its in vivo clinical applications.

### Pre-training data limitations

Gene expression typically undergoes changes following drug selection. However, the pre-trained data in this paper only consists of transcriptome data from cell lines prior to drug exposure. Due to the heterogeneity of cancer, certain cell populations may evolve into multiple resistant subpopulations under drug selection, leading to varied drug responses. Therefore, knowledge based on pre-trained GDSC data may be limited.

Furthermore, the limited availability of public datasets poses a significant challenge to the generalization capability of our model. Public datasets are often constrained in size and diversity, which can restrict their ability to fully capture the variability and complexity of real-world data. This limitation may potentially impact the model’s performance when applied to new, unseen data, as it may not have been exposed to a sufficiently broad range of scenarios and conditions during training. One potential solution is to gather a large amount of authentic clinical data for extensive training in order to enhance the model’s generalization in real-world scenarios. However, as far as we know, publicly available data remains limited. Collecting more public clinical datasets is a time-consuming and laborious task but it is crucial for improving models’ performance.

### Post-treatment transcriptome alteration

Pre-clinical models based on pretreatment transcriptome data can be applied for pre-clinical drug screening and may not able to predict drug efficacy in the subsequent stage based on post-treatment transcriptome data.

However, we have validated the capability of our model to differentiate between response concentrations of various drugs on the missing GDSC dataset and an external TCGA dataset. This demonstrates the potential of our model for conducting drug sensitivity studies based on in vitro cultured cell lines, as well as for pre-clinical drug screening.

## 6 Conclusion

This study proposes a framework for distilling privileged knowledge from a multimodal genome teacher network to a unimodal gene expression student network, resulting in an improvement of approximately 6% compared to the optimal unimodal baseline. By implementing a feature reweighting strategy, we integrate genomic knowledge into the process of extracting features from the transcriptome. This approach assists the model in focusing on crucial features. Furthermore, we utilize a relation-aware differential distillation module to further align inter-sample correlations and emphasize challenging samples. Transferable studies on clinical data further demonstrate the potential application of our model in pre-clinical drug screening. Ultimately, understanding the heterogeneity of drug responses will greatly benefit personalized oncology in the long run.

## Code and Data Availability

All datasets mentioned are publicly accessible. Genomic information: GDSC: https://www.cancerrxgene.org/gdsc1000/GDSC1000_WebResources/Home.html, CCLE: https://depmap.org/portal/data_page/?tab=allData, TCGA: https://zenodo.org/records/7264573#.Y16Ed3ZByUl. Drug response files of each platform: [GDSC (password: KDRU)](https://pan.baidu.com/s/1Z7xiX4TQyaXKydSceNhp0g), GDSC2, CCLE_PRISM, TCGA. The code will be released here https://github.com/geshuang307/KDRUG.

## Author Contributions

Shuang Ge: Writing - original draft, Conceptualization, Methodology, Writing - review & editing. Shuqing Sun: Software, Data curation. Huan Xu: Software, Visualization. Qiang Cheng: Supervision, Writing - review & editing. Zhixiang Ren: Funding acquisition, Supervision, Writing - review & editing.

## Acknowledgements

This work was supported by the Peng Cheng Laboratory Foundation under grant number PCL2021A13/004.

## Footnotes

1 https://www.cancerrxgene.org/downloads/bulk_download

2 https://zenodo.org/records/7264573#.Y16Ed3ZByUl[5]

3 https://depmap.org/portal/data_page/?tab=allData

## References

[1] Chapman, P.B., Hauschild, A., Robert, C., Haanen, J.B., Ascierto, P., Larkin, J., Dummer, R., Garbe, C., Testori, A., Maio, M., et al.: Improved survival with vemurafenib in melanoma with braf v600e mutation. New England Journal of Medicine 364(26), 2507–2516 (2011)

[2] Yang, W., Soares, J., Greninger, P., Edelman, E.J., Lightfoot, H., Forbes, S., Bindal, N., Beare, D., Smith, J.A., Thompson, I.R., et al.: Genomics of drug sensitivity in cancer (gdsc): a resource for therapeutic biomarker discovery in cancer cells. Nucleic acids research 41(D1), 955–961 (2012)

[3] Barretina, J., Caponigro, G., Stransky, N., Venkatesan, K., Margolin, A.A., Kim, S., Wilson, C.J., Lehár, J., Kryukov, G.V., Sonkin, D., et al.: The cancer cell line encyclopedia enables predictive modelling of anticancer drug sensitivity. Nature 483(7391), 603–607 (2012)

[4] Ma, J., Fong, S.H., Luo, Y., Bakkenist, C.J., Shen, J.P., Mourragui, S., Wessels, L.F., Hafner, M., Sharan, R., Peng, J., et al.: Few-shot learning creates predictive models of drug response that translate from high-throughput screens to individual patients. Nature Cancer 2(2), 233–244 (2021)

[5] Shen, B., Feng, F., Li, K., Lin, P., Ma, L., Li, H.: A systematic assessment of deep learning methods for drug response prediction: from in vitro to clinical applications. Briefings in Bioinformatics 24(1), 605 (2023)

[6] Partin, A., Brettin, T.S., Zhu, Y., Narykov, O., Clyde, A., Overbeek, J., Stevens, R.L.: Deep learning methods for drug response prediction in cancer: predominant and emerging trends. Frontiers in medicine 10, 1086097 (2023)

[7] Sharifi-Noghabi, H., Zolotareva, O., Collins, C.C., Ester, M.: Moli: multiomics late integration with deep neural networks for drug response prediction. Bioinformatics 35(14), 501–509 (2019)

[8] Ding, M.Q., Chen, L., Cooper, G.F., Young, J.D., Lu, X.: Precision oncology beyond targeted therapy: combining omics data with machine learning matches the majority of cancer cells to effective therapeutics. Molecular cancer research 16(2), 269–278 (2018)

[9] Argelaguet, R., Velten, B., Arnol, D., Dietrich, S., Zenz, T., Marioni, J.C., Buettner, F., Huber, W., Stegle, O.: Multi-omics factor analysis—a framework for unsupervised integration of multi-omics data sets. Molecular systems biology 14(6), 8124 (2018)

[10] Hu, M., Maillard, M., Zhang, Y., Ciceri, T., La Barbera, G., Bloch, I., Gori, P.: Knowledge distillation from multi-modal to mono-modal segmentation networks. In: Medical Image Computing and Computer Assisted Intervention–MICCAI 2020: 23rd International Conference, Lima, Peru, October 4–8, 2020, Proceedings, Part I 23, pp. 772–781 (2020). Springer

[11] Garcia, N.C., Morerio, P., Murino, V.: Modality distillation with multiple stream networks for action recognition. In: Proceedings of the European Conference on Computer Vision (ECCV), pp. 103–118 (2018)

[12] Tian, Y., Krishnan, D., Isola, P.: Contrastive representation distillation. arXiv preprint arXiv:1910.10699 (2019)

[13] Xing, X., Zhu, M., Chen, Z., Yuan, Y.: Comprehensive learning and adaptive teaching: Distilling multi-modal knowledge for pathological glioma grading. Medical Image Analysis 91, 102990 (2024)

[14] Costello, J.C., Heiser, L.M., Georgii, E., Gönen, M., Menden, M.P., Wang, N.J., Bansal, M., Ammad-Ud-Din, M., Hintsanen, P., Khan, S.A., et al.: A community effort to assess and improve drug sensitivity prediction algorithms. Nature biotechnology 32(12), 1202–1212 (2014)

[15] Xia, F., Allen, J., Balaprakash, P., Brettin, T., Garcia-Cardona, C., Clyde, A., Cohn, J., Doroshow, J., Duan, X., Dubinkina, V., et al.: A cross-study analysis of drug response prediction in cancer cell lines. Briefings in bioinformatics 23(1), 356 (2022)

[16] Liu, P., Li, H., Li, S., Leung, K.-S.: Improving prediction of phenotypic drug response on cancer cell lines using deep convolutional network. BMC bioinformatics 20, 1–14 (2019)

[17] Nguyen, T., Nguyen, G.T., Nguyen, T., Le, D.-H.: Graph convolutional networks for drug response prediction. IEEE/ACM transactions on computational biology and bioinformatics 19(1), 146–154 (2021)

[18] Nguyen, G.T., Vu, H.D., Le, D.-H.: Integrating molecular graph data of drugs and multiple-omic data of cell lines for drug response prediction. IEEE/ACM transactions on computational biology and bioinformatics 19(2), 710–717 (2021)

[19] Chu, T., Nguyen, T.T., Hai, B.D., Nguyen, Q.H., Nguyen, T.: Graph transformer for drug response prediction. IEEE/ACM Transactions on Computational Biology and Bioinformatics 20(2), 1065–1072 (2022)

[20] Hinton, G., Vinyals, O., Dean, J.: Distilling the knowledge in a neural network. arXiv preprint arXiv:1503.02531 (2015)

[21] Zhang, H., Chen, D., Wang, C.: Confidence-aware multi-teacher knowledge distillation. In: ICASSP 2022-2022 IEEE International Conference on Acoustics, Speech and Signal Processing (ICASSP), pp. 4498–4502 (2022). IEEE

[22] Du, S., You, S., Li, X., Wu, J., Wang, F., Qian, C., Zhang, C.: Agree to disagree: Adaptive ensemble knowledge distillation in gradient space. advances in neural information processing systems 33, 12345–12355 (2020)

[23] Yuan, F., Shou, L., Pei, J., Lin, W., Gong, M., Fu, Y., Jiang, D.: Reinforced multi-teacher selection for knowledge distillation. In: Proceedings of the AAAI Conference on Artificial Intelligence, vol. 35, pp. 14284–14291 (2021)

[24] Tomczak, K., Czerwińska, P., Wiznerowicz, M.: Review the cancer genome atlas (tcga): an immeasurable source of knowledge. Contemporary Oncology/Wspólczesna Onkologia 2015(1), 68–77 (2015)

[25] Zheng, Z., Chen, J., Chen, X., Huang, L., Xie, W., Lin, Q., Li, X., Wong, K.-C.: Enabling single-cell drug response annotations from bulk rna-seq using scad. Advanced Science 10(11), 2204113 (2023)

[26] Corsello, S.M., Nagari, R.T., Spangler, R.D., Rossen, J., Kocak, M., Bryan, J.G., Humeidi, R., Peck, D., Wu, X., Tang, A.A., et al.: Non-oncology drugs are a source of previously unappreciated anti-cancer activity. BioRxiv, 730119 (2019)

[27] Schölkopf, B., Smola, A., Müller, K.-R.: Kernel principal component analysis. In: International Conference on Artificial Neural Networks, pp. 583–588 (1997). Springer

[28] Weininger, D.: Smiles, a chemical language and information system. 1. introduction to methodology and encoding rules. Journal of chemical information and computer sciences 28(1), 31–36 (1988)

[29] Scarselli, F., Gori, M., Tsoi, A.C., Hagenbuchner, M., Monfardini, G.: The graph neural network model. IEEE transactions on neural networks 20(1), 61–80 (2008)

[30] Vapnik, V., Vashist, A.: A new learning paradigm: Learning using privileged information. Neural networks 22(5-6), 544–557 (2009)

[31] Garcia, N.C., Morerio, P., Murino, V.: Learning with privileged information via adversarial discriminative modality distillation. IEEE transactions on pattern analysis and machine intelligence 42(10), 2581–2593 (2019)

[32] Liu, Y., Fan, Q., Zhang, S., Dong, H., Funkhouser, T., Yi, L.: Contrastive multimodal fusion with tupleinfonce. In: Proceedings of the IEEE/CVF International Conference on Computer Vision, pp. 754–763 (2021)

[33] Yang, J., Martinez, B., Bulat, A., Tzimiropoulos, G., et al.: Knowledge distillation via softmax regression representation learning. (2021). International Conference on Learning Representations (ICLR)

[34] Vaswani, A., Shazeer, N., Parmar, N., Uszkoreit, J., Jones, L., Gomez, A.N., 22 Kaiser, L., Polosukhin, I.: Attention is all you need. Advances in neural information processing systems 30 (2017)

[35] Zhou, X., Jiang, Z., Okuwobi, I.P.: Cafnet: Cross-attention fusion network for infrared and low illumination visible-light image. Neural Processing Letters 55(5), 6027–6041 (2023)

[36] Li, G., Li, X., Wang, Y., Zhang, S., Wu, Y., Liang, D.: Knowledge distillation for object detection via rank mimicking and prediction-guided feature imitation. In: Proceedings of the AAAI Conference on Artificial Intelligence, vol. 36, pp. 1306–1313 (2022)

[37] Paszke, A., Gross, S., Massa, F., Lerer, A., Bradbury, J., Chanan, G., Killeen, T., Lin, Z., Gimelshein, N., Antiga, L., et al.: Pytorch: An imperative style, highperformance deep learning library. Advances in neural information processing systems 32 (2019)

[38] Jiang, L., Jiang, C., Yu, X., Fu, R., Jin, S., Liu, X.: Deeptta: a transformer-based model for predicting cancer drug response. Briefings in bioinformatics 23(3), 100 (2022)

[39] Fisher, R.I., Bernstein, S.H., Kahl, B.S., Djulbegovic, B., Robertson, M.J., De Vos, S., Epner, E., Krishnan, A., Leonard, J.P., Lonial, S., et al.: Multicenter phase ii study of bortezomib in patients with relapsed or refractory mantle cell lymphoma. Journal of clinical oncology 24(30), 4867–4874 (2006)

[40] Richardson, P.G., Barlogie, B., Berenson, J., Singhal, S., Jagannath, S., Irwin, D., Rajkumar, S.V., Srkalovic, G., Alsina, M., Alexanian, R., et al.: A phase 2 study of bortezomib in relapsed, refractory myeloma. New England Journal of Medicine 348(26), 2609–2617 (2003)

[41] Huang, Z., Wu, Y., Zhou, X., Xu, J., Zhu, W., Shu, Y., Liu, P.: Efficacy of therapy with bortezomib in solid tumors: a review based on 32 clinical trials. Future oncology 10(10), 1795–1807 (2014)

[42] Tseng, L.-M., Liu, C.-Y., Chang, K.-C., Chu, P.-Y., Shiau, C.-W., Chen, K.-F.: Cip2a is a target of bortezomib in human triple negative breast cancer cells. Breast cancer research 14, 1–14 (2012)

[43] Cortes, J., Thomas, D., Koller, C., Giles, F., Estey, E., Faderl, S., Garcia-Manero, G., McConkey, D., Patel, G., Guerciolini, R., et al.: Phase i study of bortezomib in refractory or relapsed acute leukemias. Clinical Cancer Research 10(10), 3371–3376 (2004)

